# Meta-analysis of gene activity (MAGA) contributions and correlation with gene expression, through GAGAM

**DOI:** 10.1101/2023.04.04.535522

**Authors:** Lorenzo Martini, Roberta Bardini, Alessandro Savino, Stefano Di Carlo

## Abstract

It is well-known how sequencing technologies propelled cellular biology research in the latest years, giving an incredible insight into the basic mechanisms of cells. Single-cell RNA sequencing is at the front in this field, with Single-cell ATAC sequencing supporting it and becoming more popular. In this regard, multi-modal technologies play a crucial role, allowing the possibility to perform the mentioned sequencing modalities simultaneously on the same cells. Yet, there still needs to be a clear and dedicated way to analyze this multi-modal data. One of the current methods is to calculate the Gene Activity Matrix, which summarizes the accessibility of the genes at the genomic level, to have a more direct link with the transcriptomic data. However, this concept is not well-defined, and it is unclear how various accessible regions impact the expression of the genes. Therefore, this work presents a meta-analysis of the Gene Activity matrix based on the Genomic-Annotated Gene Activity Matrix model, aiming to investigate the different influences of its contributions on the activity and their correlation with the expression. This allows having a better grasp on how the different functional regions of the genome affect not only the activity but also the expression of the genes.

## 1 Introduction

Next Generation Sequencing (NGS) technologies are the backbone of the latest cellular biology research. With their incredible power to investigate fundamental cell mechanisms, NGS technologies enable the study of cellular states with high resolution, which is crucial to investigate cellular heterogeneity.

The single-cell RNA sequencing (scRNA-seq) technology is the most widely employed technology to study thousands of single-cell transcriptional profiles and investigate cellular heterogeneity based on gene expression [4]. In addition, single-cell assays for transposase-accessible chromatin sequencing (scATAC-seq) is becoming popular. Thanks to its ability to probe the whole genome and assess the accessible chromatin regions, scATAC-seq provides a complementary insight into the fundamental process of gene regulation [3] and expression [2].

These two faces of the same medal give an unprecedented way to investigate these complex mechanisms through their joint analysis. So, it is not surprising that multi-modal technologies, which allow simultaneously assessing both scRNA-seq and scATAC-seq from the same cells, are becoming crucial when investigating cell-related phenomena, including heterogeneity [6]. However, the intrinsic difference in data type between the two technologies poses some caveats to a proper joint analysis [9] [19].

In general, it is not trivial to correlate the accessibility of a particular region of the genome to gene expression, given the incredible and complex machinery involved in gene regulation. This means that scATAC-seq datasets are built considering genes as prominent features. In contrast, scATAC-seq datasets consider genomic regions as features, making integrating these two data difficult.

The concept of Gene Activity (GA) is a viable approach to correlate accessibility with gene expression [17]. The GA summarizes the genomic accessibility information in a form where the features are genes instead of genomic regions, representing how much the gene is accessible and potentially transcribed. It enables the translation of scATAC-seq data into a matrix formally similar and directly comparable to the scRNA-seq matrix, allowing for a direct investigation of the correlation between the two biological levels. However, no clear definition exists of how to model the relationship between accessible regions and genes.

A promising approach to solve this problem is the Genomic-Annotated Gene Activity Matrix (GAGAM) approach [15] [16]. In GAGAM, the association between genomic regions and accessible genes relies on a genomic model based on genomic annotations. This model constructs a Gene Activity Matrix (GAM) consisting of three different contributions associated with different functional genomic regions (i.e., promoters, exons, and enhancers). This should better model the gene regulatory landscape crucial to understanding gene expression, not just the simple gene body accessibility. GAGAM has, therefore, the peculiarity of creating a simple model that integrates different scATAC-seq signals, thus understanding and treating differently the functional information related to the accessible regions. However, in GAGAM, the complexity of the gene regulation mechanism remains hidden, and the relationship and interaction between specific regulatory elements and the gene bodies are still not represented, and, more specifically, how their accessibility impacts the actual expression remains implicit. This modeling limitation consequentially restrains the ability of GAGAM to represent the entire gene regulation mechanism accurately.

This work presents a complete meta-analysis of the GA contributions of GAGAM, starting from a multi-modal dataset to pave the way for more effective analysis. The analysis uses these data to understand better the correlation between GA and expression. Results presented in this paper help improve the definition of GA models, accurately represent the role of gene regulatory mechanisms, and support the investigation of the complex relations between DNA accessibility and gene expression. Eventually, it will also help the single-cell study of rising multi-modal datasets.

## 2 Background

### 2.1 Single-cell sequencing technologies

To fully understand the proposed analysis, it is crucial to introduce the basic technologies involved in this work. First, a quick explanation of the scATAC-seq data helps understand the derived concept of GA. scATAC-seq is a technology to provide information on the epigenomic state of the cells by probing the whole genome, which leverages the Tn5-transposase to detect all the regions where the chromatin is open, and the DNA sequence is accessible [20]. Through that, it is possible to investigate not only the genes (as for scRNA-seq) but also various functional elements, like enhancers and promoters, [11], that are scattered all over the genome but are crucial for gene regulation [7]. While scRNA-seq data have genes as features, scATAC-seq data use peaks, i.e., short genomic regions described by their coordinates on the chromosomes. This intrinsic difference poses a considerable hurdle when correlating the two biological levels. One way to overcome this is to transform the peaks into gene-like data and compare the two technologies. As mentioned in section 1, GA is one way to do so [6].

However, the current models to define GA tend to oversimplify the relationship between a gene and the accessibility of its genomic region. Specifically, some approaches like GeneScoring [13] and Signac [18] look indiscriminately to the peak signals overlapping the gene body regions without distinction between coding and non-coding regulatory elements. On the other hand, Cicero [6] defines its GA in a more structured way, considering other regulatory regions but collapsing the gene region to a single base. These methods generally retain little biological information from the raw scATAC-seq data, usually only related to gene coding regions, even if they represent only a tiny percentage of the whole signal [5]. Beyond these simplistic models, other approaches aim to comprise more accessible genomic regions and their impact on overall GA. Specifically, this work employs GAGAM since it uses curated genomic annotations to functionally label the peaks and, consequentially, associates them with the genes through a simple model [15].

### 2.2 GAGAM

GAGAM comprises information on several DNA regions, particularly from exons and non-coding regions with a regulatory role (i.e., the promoter and the enhancers of genes). This represents the main strength of GAGAM, which tries to analyze this biological information from scATAC-seq data and put it together in a model-driven method, more representative of the biological level, and could better support the cellular heterogeneity study. Therefore, GAGAM poses the basis for a more detailed investigation of the relationship between accessibility and expression in single-cell data. Its modular structure lets us control which contributions to consider in the analysis and study their relation to expression levels individually. The latter is important, especially when considering the role of regulatory regions, whose relation with gene expression is challenging to investigate. Systematically analyzing their accessibility to gene expression could build new ways to study the complex matter of gene regulation. For this reason, this work proposes a meta-analysis of the general relationship between promoter, exon, and enhancer contributions and their prevalence in the GAGAM model and a study of the correlation of each of them with gene expression.

Let us briefly recall the workflow required to construct GAGAM, starting from the raw data, to provide the reader with the necessary background. One everyday use of single-cell data is to study cellular heterogeneity, that is, to highlight the key changing features that characterize the different types of cells. GAGAM also goes in this direction. All methods to analyze single-cell data require properly preprocessing the initial data. Indeed, before constructing GAGAM itself, one preprocesses both parts of the multi-modal data (i.e., the scRNA-seq and scATAC-seq matrices). This preprocessing includes (i) Quality Control check, (ii) normalization and standardization, (iii) Principal Component Analysis (PCA), (iv) Uniform Manifold Approximation and Projection (UMAP) dimensionality reduction, (v) clustering, and (vi) Differential Expression (DE) analysis [14]. In particular, clustering and DE are the most relevant results, providing a reliable figure of the dataset’s cell states/cell types, giving ground for the following investigations.

GAGAM, works on the preprocessed scATAC-seq data alone, organized in the form of a matrix **D**_|*P*|×|*C*|_, where *P* is the set peaks in the detaset, and *C* is the set of avilable cells. As shown in Algorithm 1 (lines 1 to 5), first, GAGAM employs the UCSC Genome Browser [12] to obtain genomic annotations and label all the peaks *p* ∈ *P* overlapping with regions of interest (i.e., promoters, genes’ exons, and enhancers), assigning to them the respective labels `prom, exon, enhD`. In this way, it is possible to understand the function of the accessible peaks and, consequentially, relate the function to the genes. Then it constructs three label-specific matrices (Algorithm 1, lines 6 to 8), which constitute the three contributions investigated in this work, denoted as 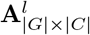 where *G* is the set of genes with at least one peak mapping to them and *l* ∈ {*prom, enhd, exon*}. Finally, GAGAM sums all the contributions weighted by model-specific weights to obtain the final GA matrix. In the GAGAM implementation introduced in [15], simple binary weights are used, but a better understanding of the contribution of the three matrices could help fine-tune them. The final model of GAGAM activity translates the simple accessibility data from scATAC-seq into a score representing the overall accessibility of the gene and its potential to be expressed. This simple structure allows for investigating each contribution individually in a direct way, which is the core of this work and a vital element of the potentiality of GAGAM itself.

#### Algorithm 1 GAGAM construction

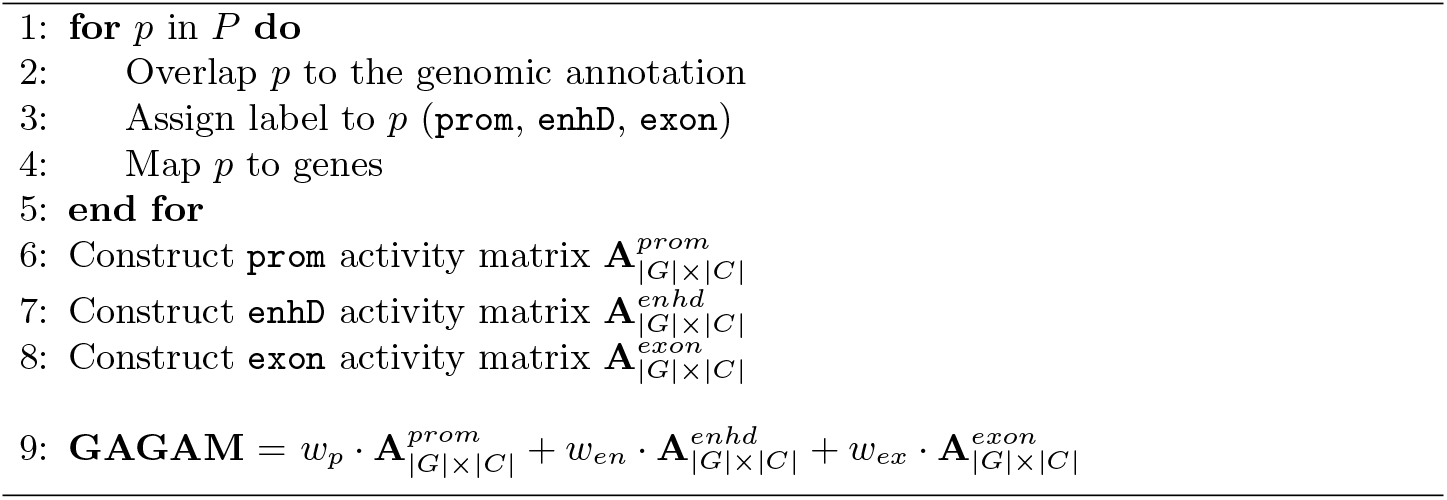

## 3 Meta-Analysis

A multi-modal dataset is required to jointly analyze Gene Activity *per se* and gene expression. The dataset of choice is an open-access dataset from the 10X Genomics platform, consisting of 10,691 cells from adult murine peripheral blood mononuclear cell (PBMC) [1]. The scATAC-seq part of the dataset has a total of 115,179 peaks as features, while the scRNA-seq part has 36,601 genes. The tools employed to process and elaborate the data are GAGAM (the focus of this paper, accessible from [15]) and Seurat [4]. The latter is one of the most well-known and highly-utilized single-cell pipelines. All the code employed for this work is available at https://github.com/smilies-polito/MAGA, including all the supplementary material and figures.

Before starting with the actual meta-analysis, it is worth noting that scRNA-seq and scATAC-seq detect only a tiny fraction of the actual signal from each cell (around 10-45% for scRNA-seq and only 1-10% for scATAC-seq [5]). This translates into considerable sparsity for the data. For each cell, the dataset contains several zero entries that could be false negatives [10]. This characteristic introduces noise when trying to correlate accessibility and expression. For this reason, this work explores the idea of performing the analysis based on the concept of *aggregated cell* behavior. Specifically, it aggregates cells from the same clusters obtained from preprocessing the raw scRNA-seq data, representing the average over groups of cells instead of single cells. This way, these clusters should, with high probability, represent the cell types [4]; thus, exploring them could be relevant for cellular heterogeneity studies.

The procedure to calculate gene activity and expression of the aggregated cells is described in Algorithm 2. For the subsequent analyses, let us denote with **A**_|*G*|×|*AC*|_ the aggregated cells activity matrix and with **E**_|*G*|×|*AC*|_ the aggregated cells expression matrix, where *G* is the set of genes and *AC* is the set of aggregated cells.

### Algorithm 2 Aggregated cells definition

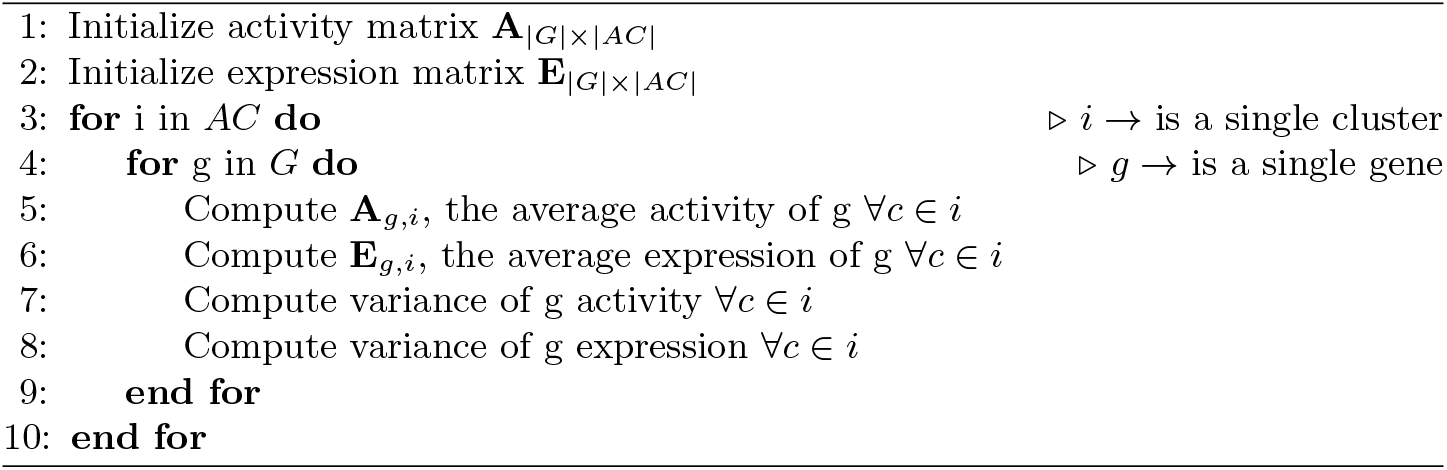

The meta-analysis focuses on three separate investigations reported in the following subsections.

### 3.1 Peaks Information

After processing the data and obtaining the GAGAM contributions, the first investigation focuses on the three labels (i.e., `prom, exon, enhD`), assigned to peaks during the GAGAM construction (Algorithm 1 line 3), and the information they carry on. Indeed, understanding what type of information and how much is retained from the raw scATAC-seq data is crucial for creating models from them. As discussed in Section 2, while other GA methods look into gene regions only, GAGAM focuses on more regions of the genome. This type of analysis supports the correctness of this choice. Moreover, showing the non-equal distribution of the labels justifies the separate investigation of the three contributions performed in the following sections.

Let us focus on the proportions between non-labeled, prom, exon, and enhancers labels. This proportion gives direct knowledge of how much information GAGAM retains from the raw scATAC-seq, which is not trivial [20]. Furthermore, it is relevant to investigate the peak-to-gene assignments. From GAGAM computation, it is also possible to retrieve the link between peaks and genes; thus, it is straightforward to study how many and which labeled peaks the model assigns to the genes. This simple analysis gives insight into the general GA model constitution, investigating how much accessibility information relates to each gene, which is crucial for the improvement of the model itself.

Figure 1 shows some information about peak labeling. Of the 115,179 peaks, 92,100 (80%) received one of the labels. GAGAM does not necessarily employ all labeled peaks since it filters out the ones that do not associate with a gene (namely, for some promoter and enhancer peaks.). Among the labeled peaks, there is a clear predominance of the enhancer peaks (about 34% of all peaks) against a limited portion of promoter peaks (about 13% of all peaks), remarking that the epigenetic information from the scATAC-seq data comes from distal regions and not just near the gene coding regions.

**Fig. 1.**
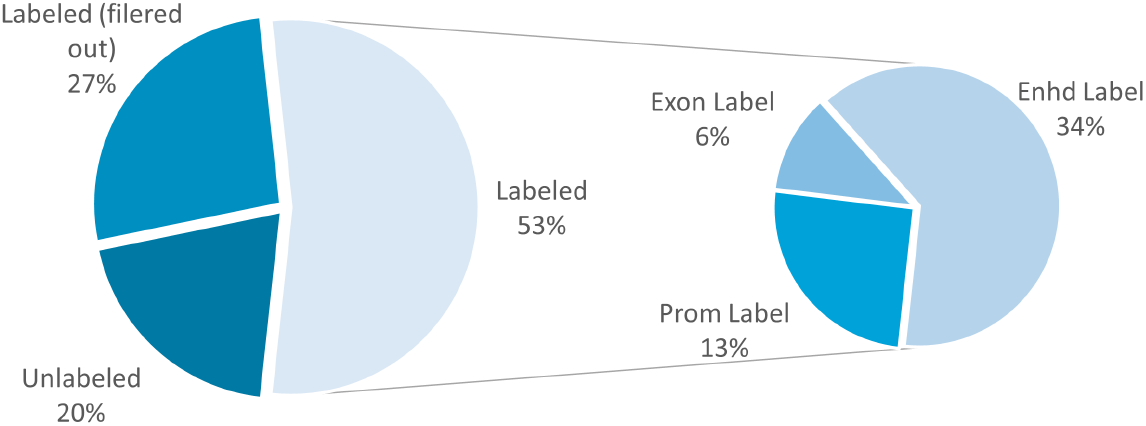
The figure shows the distribution of labels among the peaks. Additionally, the labeled peaks are divided into three different labels. Most peaks receive a label, even though some data is not considered. Most labels are enhancers, showing how much of the information and potentiality of the scATAC-seq data comes from non-gene-related regions.

Figure 2 presents the number of labeled peaks assigned to genes for each label type. The vast majority of genes (86%) have only one promoter peak set to them, which appears to be the average case. However, a few genes have multiple promoter peaks mapping to them, which is entirely unexpected. Directly examining the UCSC genome browser, it becomes clear that the multiple promoters map to different isoforms of the same genes. This shows the capability of GAGAM to have detailed resolution on the GA, meaning it can independently explore several isoforms of genes.

**Fig. 2.**
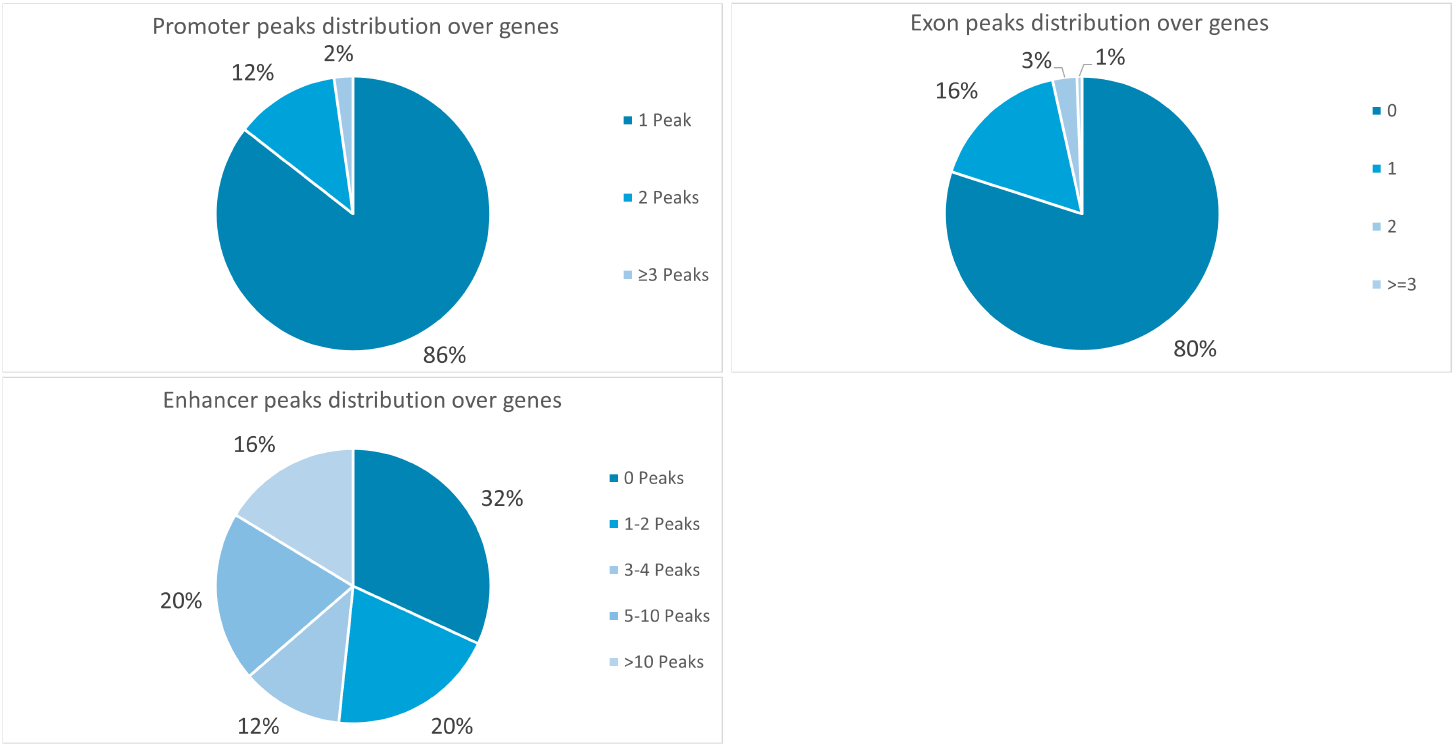
Distribution of the number of peaks mapping to genes, divided by label types. The percentages represent the percentage of all genes with n peaks mapping to them.

Regarding the exon peaks, most genes do not have any mapping to them. One of the reasons stems from the small number of exon peaks (namely, 7046, only 6% of the total peaks), so they only cover some genes.

On the other hand, the genes tend to have many enhancer peaks linked to them, also because, unlike the other labeled peaks, a single enhancer can map to multiple genes. The latter highlights the relevance of creating a reliable model to connect them to genes, which is central to future developments.

### 3.2 Activity-Expression correlation

Our previous work [15] qualitatively addressed this type of investigation to broadly study Activity-Expression (A-E) patterns for specific genes. This paper applies a quantitative approach. Pearson coefficient [8] computed between the activity and expression of each gene on aggregated cells is used to quantify the A-E correlation, and this information is visualized through a set A-E plots. The investigation is performed independently on each peak label from the previous section. This enables us to investigate how each accessibility label affects the gene expression and better tune the GA models.

The promoter peaks are expected to be most correlated with the expression [20,18]. The correlation is relatively high for most genes, with 71% correlating higher than 0.5 with statistically significant p-values (*p* < 0.05). Moreover, focusing on the top 1,000 genes with the lowest variance, 94% of them correlate higher than 0.5, strengthening the previous result. It is also interesting to notice that considering the list of DE genes, the correlation is still relevant, with 91% being over the mentioned threshold. Therefore, it is fair to state that the accessibility of the gene’s promoter results in its detectable and subsequent expression.

Besides a purely numerical analysis, it is relevant to visualize the correlation. For each aggregated cell, it is possible to plot each gene as a point in the A-E space and explore the general trend of the genes inside the clusters. This way, one can investigate the difference between clusters that could convey pertinent information for the cellular heterogeneity study. For the sake of space and clarity, only the first four A-E plots (denoted as CL0, CL1, CL2, and CL3) are reported in Figure 3. Information written in these plots is representative of the overall results. All remaining plots and figures are available at https://github.com/smilies-polito/MAGA. The points are on a log space to allow better visualization, while the colors represent meaningful information on each gene. The black points are all the genes, while the red points represent the differentially expressed genes for the cluster. The latter comes from the DE analysis performed on the clusters, precisely the top specific marker genes per cluster. Moreover, the plots contain green dashed lines defining the genes’ mean activity or expression in that specific aggregated cell.

**Fig. 3.**
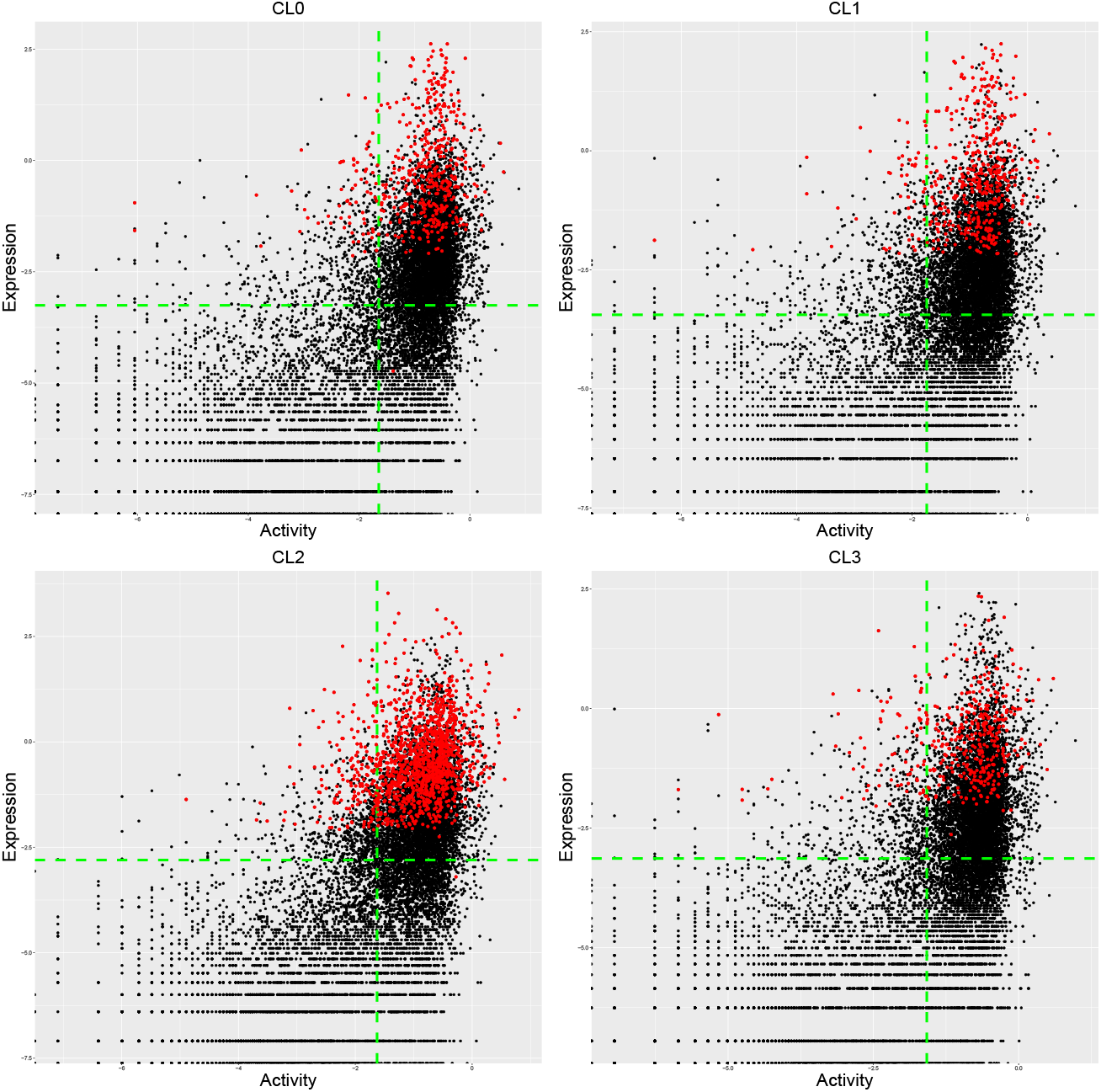
Activity-Expression plots. Each point represents a gene, with its expression and promoter activity from the aggregated cells. Here are present only the first four aggregated cells. The red dots represent the DE genes from the cluster

Figure 3 shows that most points are in the top-right area, meaning high expression and activity, demonstrating the correlation between the two characteristics. Most of the DE genes from that specific cluster (points in red) lie in this area, highlighting their relevant high activity other than expression. The least populated area is the top-left area, representing the low activity high expression area, while the opposite is pretty populated. This observation is not trivial, and it highlights that the promoter accessibility of a gene can implicate various levels of expression. However, when the accessibility is low or null, the genes are rarely expressed.

The exon contribution is the smallest. Only 7,046 genes have at least one exon peak linked to them. Nonetheless, their specific information can be informative to the model. The Pearson correlation on the aggregated cells reveals that only about 55% of the remaining genes correlate greater than 0.5, with the percentage lowering to about 41% when considering the lowest variance genes. This is more clear from the A-E plots (Figure 4), where one can see that the points tend to occupy the right-most part consistently, indicating the high activity area. Differently from the promoter activity, which was spread out and ranged on different levels, the exon activity appears more set in a binary-like fashion. The exon contribution to the activity seems to have a relatively high contribution or not to be present at all. This also explains the lack of a high correlation with the expression since the exon contribution does not appear to influence it strongly.

**Fig. 4.**
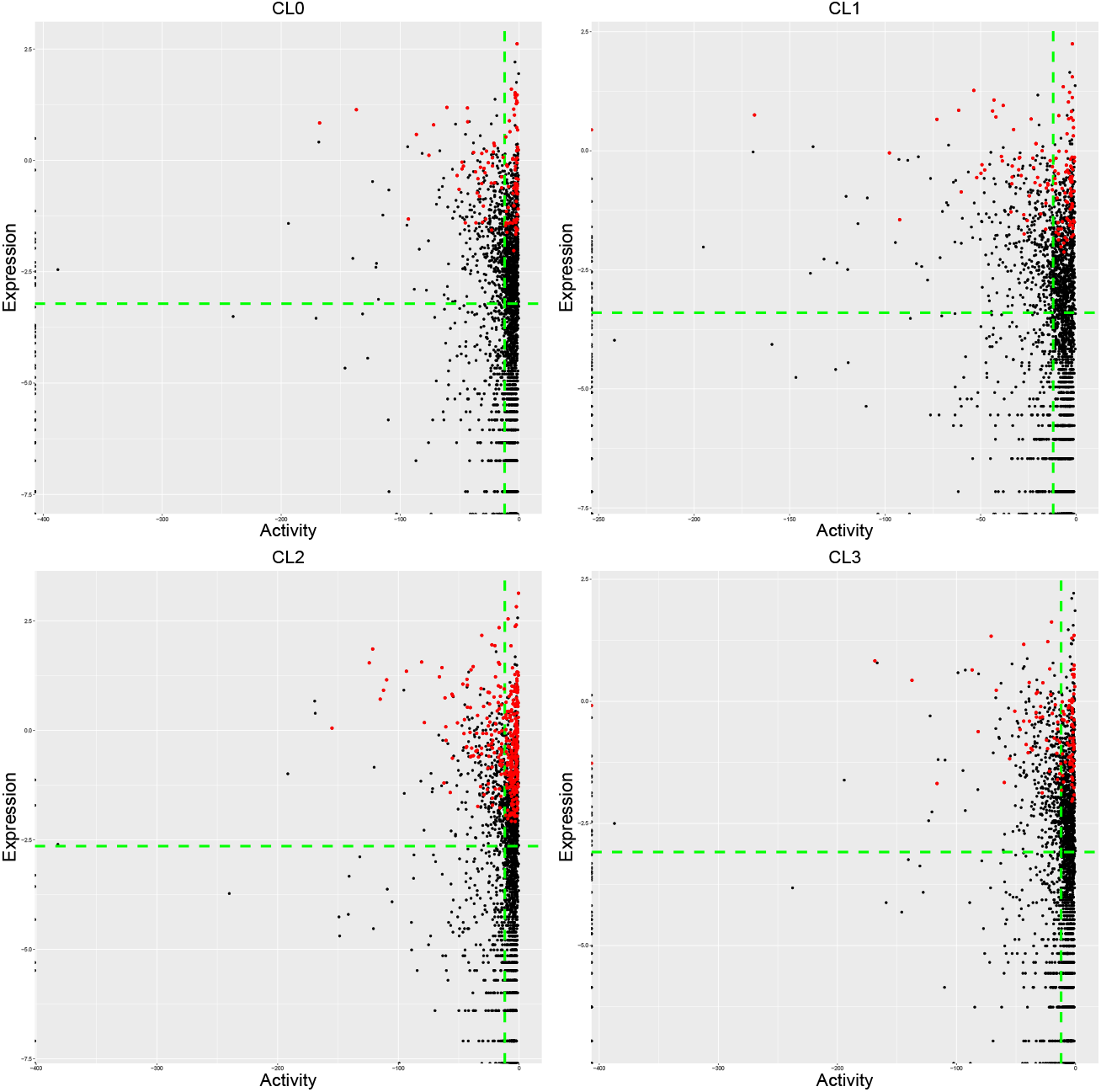
A-E plots. Each point represents a gene, with its expression and exon activity from the aggregated cells. Here are present only the first four aggregated cells. The red dots represent the DE genes from the cluster.

The enhancer contribution is the most complex and potentially important to explain gene regulation. Many different enhancer peaks can map to a gene, and each can map to various other genes, making the correlation less trivial to study. The Pearson correlation confirms that about 63% of the genes correlate with enhancer activity and expression higher than the threshold of 0.5. It is lower than the promoter contribution but higher than the exon one. When considering the lowest variable genes, differently from the exon case, this percentage rises to 70%. Eventually, the correlation calculated for the DE genes improves, with 72% of them being over the threshold.

The results in Figure 5 differ slightly from promoters and exons. The points still predominate in the top-right area but spread out more, showing that the correlation is less predominant. This is prominent in specific clusters like the “CL2”, where many genes have discordant activity and expression, meaning that the enhancer activity contribution is less necessary for the expression than the other contributions. The red points, representing the DE genes, still reside in the right-top area, even if more to the top-left.

**Fig. 5.**
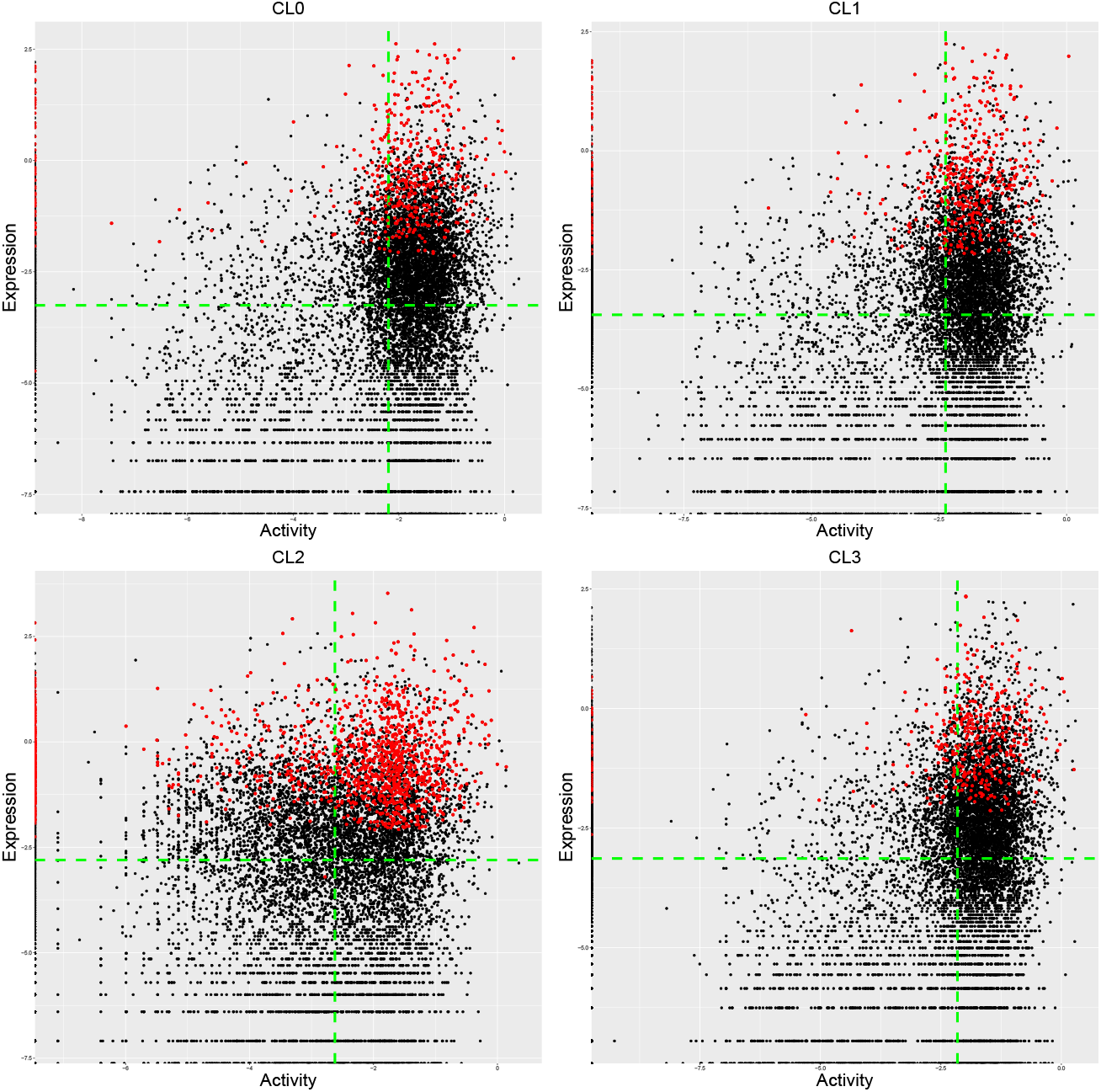
A-E plots. Each point represents a gene, with its expression and enhancer activity from the aggregated cells. Here are present only the first four cluster aggregated cells. The red dots represent the DE genes from the cluster

Intriguingly, the Pearson correlation on the DE genes remains relatively high for all peak labels. This can be ascribed to the fact that these genes, which have, by definition, high expression variability among the clusters, must also show a similar activity variability at all levels. Therefore, the concept of marker genes, well-known in transcriptomic analysis studies, could even be applied to epigenomic studies. However, it is curious that sometimes these genes are expressed despite a low or even null activity, as in the enhancer case. At first glance, it might appear counterintuitive to have expression and no GA, which could stem from the noise caused by the different depths of the technologies. However, the epigenetic and transcript levels work on very different regulation time scales, meaning a change in accessibility does not immediately propagates to the expression. Therefore, the case could represent an informative dynamical change undetected from the static view given by the separated data. More investigations will be needed to confirm such a hypothesis.

### 3.3 Activity-Expression Coherence

Section 3.2 discussed the general correlation between activity and expression. Yet, there are some cases where the causal relationship between them could be more complex (e.g., for the enhancer contribution). Therefore, it is intriguing to investigate the coherence between expression and activity, meaning how much the presence of activity of genes implicates their expression in a binary way. This analysis studies how many genes with an activity greater than zero also have an expression greater than zero and vice-versa (see Algorithm 3). The algorithm reduces the activity *(a)* and expression (*e*) in the **A**_|*G*|×|*AC*|_ and **E**_|*G*|×|*AC*|_ matrices to a binary form (lines 1-14), considering positive values as one while negative or null values as 0. Then it counts genes falling under four cases (lines 15-31) based on the combination of *a* and *e* of each gene: (i) high activity and expression (line 19), (ii) high activity but low expression (line 21), (iii) low activity but high expression (line 25), and (iv) low activity and expression.

#### Algorithm 3 Coherence calculation

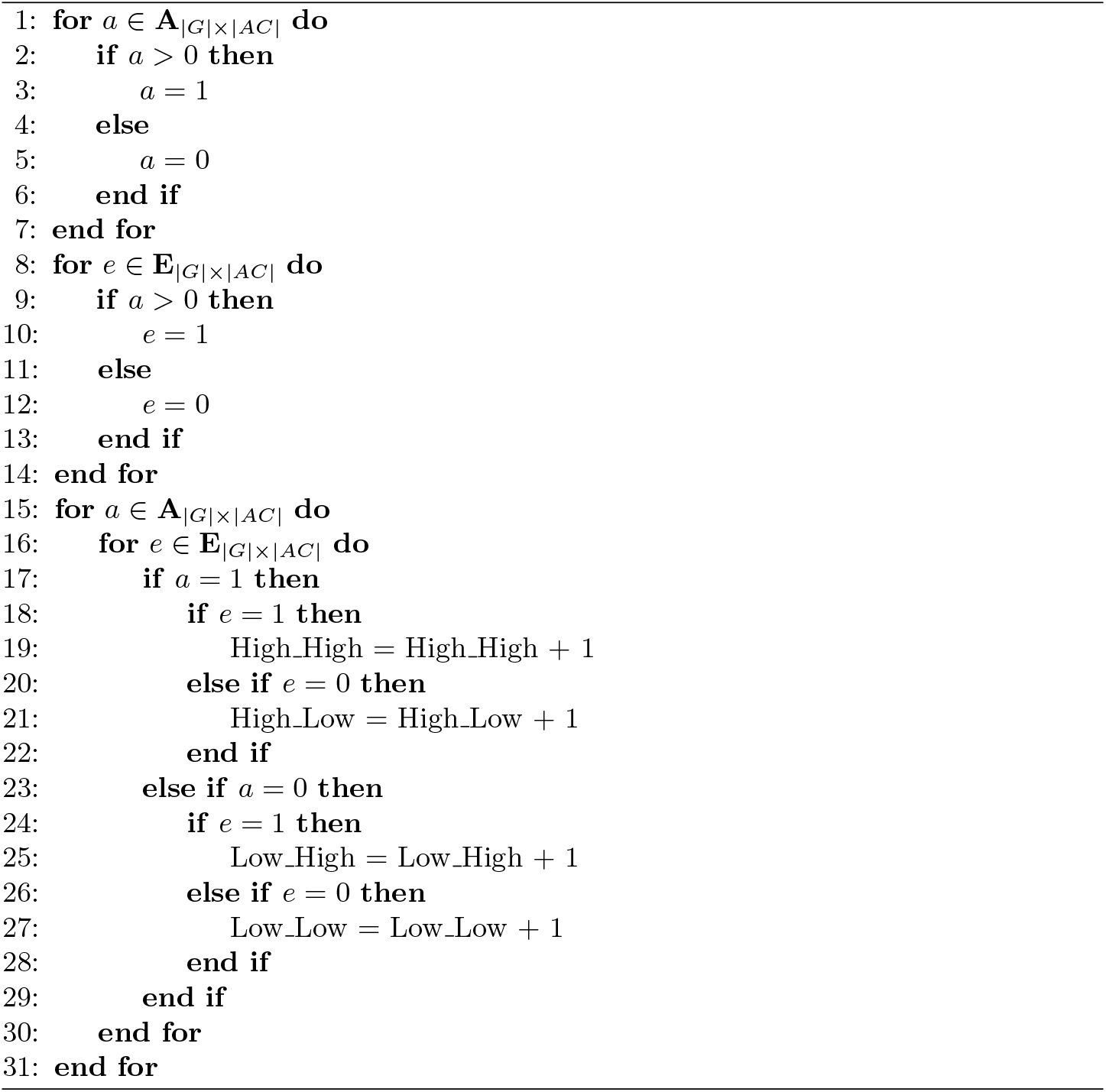

This approach highlights the interesting non-trivial case where activity and expression exhibit low or null coherence. Table 1 reports all the results as the average percentage of genes in the four cases, distinguished by peak label.

**Table 1.**
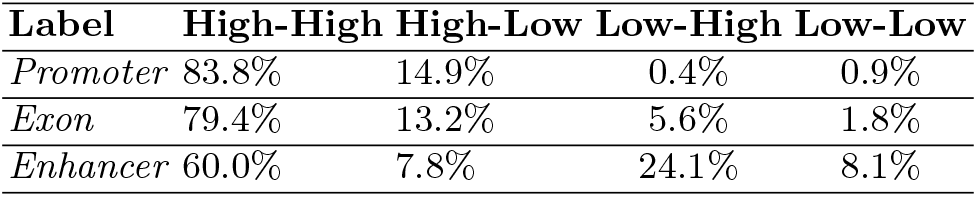
Activity-Expression coherence: The table shows the percentages of genes in the four cases. Namely, they are High activity High expression, High activity Low expression, Low activity High expression, and Low activity Low expression.

The A-E coherence for the promoter contribution is in line with the correlation results as about 83% belong to the high-high class and about 15% to the high-low case. It remarks the fact that promoter accessibility is necessary but not sufficient for the expression.

Unlike the correlation results, in the case of exons, the high-high case still includes most of the genes, despite being in a lower percentage than previously. It is relevant to notice the increase in the low-high case, which could support the hypothesis of the possible dynamical transcription changes. Indeed, the exon accessibility in gene regulation could follow different timings than promoters and be a better probe for these dynamical changes.

Finally, the A-E coherence is the most surprising for the enhancer contribution. The high-high count in the enhancer case lowers to 60% in favor of a significant increase in the low-high case. This differs from the 80% of all genes identified during correlation analysis. The low-high case was almost nonexistent when considering correlation, but now it includes about 24% of the genes. Therefore, the enhancer activity seems not a strictly necessary condition for the expression, although, when present, it positively correlates with it. Many reasons may justify this behavior. The modeling of enhancers on gene regulation is far from trivial, and for clarity, the GAGAM model simplifies it. As previously mentioned, one enhancer peak can map to many genes and influence their activity simultaneously. At the same time, in a single cell, it is likely to impact only a subset of them at a time. Moreover, the current model cannot distinguish between enhancers and silencers, which affects the sign of the contribution to the expression. Finally, the timescales involving the enhancers in gene regulation are probably even more dilated than the previous contributions. The low-high correlation could represent a dynamic change in the gene expression, as mentioned before. Despite these limitations, the results show the model can retrieve a significant correlation between enhancer contribution and expression.

In general, the A-E coherence results emphasize that the accessibility of a gene is not a sufficient condition for the expression, and in some cases, neither is necessary. The reason behind this not-trivial behavior could be different and exciting to investigate. Specifically, the enhancer contribution seems the most complex but potentially informative for gene regulation, and its understanding and fine-tuned interpretation could unravel crucial details. However, this is not part of this work and is left for future development.

### 3.4 Conclusions

This work presented a meta-analysis of GAGAM informative content and, precisely, how its building blocks correlate with the expression on a multimodal single-cell dataset. The results are pretty revealing. First, from the peak information analysis (Section 3.1), one can immediately understand that the information retained by GA from the raw scATAC-seq data is limited and diversified. Indeed, only about half of the original peaks are considered, with them being unevenly distributed between the three labels. In particular, there is an evident predominance of enhancer peaks, which highlights the limitations of other GA methods (Section 2) since they focus only on promoter and gene body regions, which cover only a tiny portion of the epigenetic data information. Understanding that an optimal GAM should retain as much information from the raw data as possible is fundamental, especially for accurately modeling the epigenetic level.

Regarding the A-E correlation, this meta-analysis focused on studying how the three contributions of GAGAM correlate with the actual expression. First, the promoter contribution shows the most linear behavior, meaning that genes with active promoters also consistently display expression. On the other hand, the exon contribution already has less clear and trivial results; namely, the exon accessibility has a remarkably lower correlation with the expression than the other two. Lastly, the enhancers show the most complex results. In particular, the A-E plots and the A-E coherence highlights the significant number of discordant genes (i.e., the genes with High activity Low expression and Low activity High expression), which emphasizes the intricate relationship between gene expression and activity.

In general, the incoherencies between gene activity and expression are interesting. Indeed, the different time scales at which transcriptomic and epigenomics work could cause these discrepancies, giving insight into the dynamic changes that are part of gene regulation. This fact highlights the power and relevance of studying multimodal data through the GAM, which could help go beyond the intrinsic static nature of single-cell data.

In any case, this analysis is crucial to improving the underlying GA model. Comprehending the relationship between specific genomic regions’ activity and gene expression can help fine-tune each contribution’s weights on the final matrix. Moreover, better models also go toward a more accurate representation of the gene regulatory mechanism, opening the possibility of investigating in new ways. Lastly, this type of meta-analysis can become an extra tool for studying the increasingly popular multimodal datasets and help the joint analysis of scRNA-seq and scATAC-seq.

